# Interaction of thyroid hormones and gonadotropin inhibitory hormone in the multifactorial control of zebrafish (*Danio rerio*) spermatogenesis

**DOI:** 10.1101/2021.03.01.433429

**Authors:** Maira S. Rodrigues, Hamideh P. Fallah, Maya Zanardini, Hamid R. Habibi, Rafael H. Nóbrega

## Abstract

Reproduction is under multifactorial control of neurohormones, pituitary gonadotropins, as well as a number of gonadal hormones including sex steroids and growth factors. Gonadotropin-inhibitory hormone (Gnih), a novel RFamide neuropeptide, was shown to be involved in the control of pituitary gonadotropin production, as well as being involved as a paracrine factor in the regulation of gonadal function. In this context, recent studies have demonstrated that Gnih inhibited gonadotropin-induced spermatogenesis in the zebrafish testicular explants. Thyroid hormones are known to interact with the reproductive axis, and are, in particular, involved in the regulation of testicular function. Based on this background, we investigated the interaction between Gnih and thyroid hormones in the control of zebrafish spermatogenesis. To this end, zebrafish adult males were treated with the goitrogen methimazole (1mM for 21 days) in order to generate a hypothyroid model organism. Subsequently, a factorial design using an *ex vivo* testis culture system in combination with histomorphometrical and FACScan cell cycle analyses were adopted. Our results showed that methimazole treatment affected both basal and gonadotropin-induced spermatogenesis, in particular, meiosis and spermiogenesis. Moreover, the goitrogen treatment nullified the inhibitory actions of Gnih on the gonadotropin-induced spermatogenesis, specifically in the haploid cell population. We have demonstrated that thyroid hormones interaction with gonadotropin and Gnih are important components for the regulation of zebrafish spermatogenesis. The results provide a support for the hypothesis that thyroid hormones are important contributors in multifactorial control of spermatogenesis in zebrafish.

## 1. Introduction

In vertebrates, reproduction is under control of multiple signals from the brain-pituitary-gonadal axis (Habibi and Andreu-Vieyra, 2007; Nóbrega et al., 2010; Schulz et al., 2010; Tsutsui et al., 2010; Habibi et al., 2012; Tsutsui et al., 2013; Tovo-Neto et al., 2018; Tovo-Neto et al., 2020; Ma et al., 2020 a, b). The gonadotropin-releasing hormone (Gnrh) is one of the key players of the multifactorial regulation of reproduction due to its central role on stimulating the synthesis and secretion of the gonadotropin hormones. The gonadotropins, follicle-stimulating hormone (Fsh) and luteinizing hormone (Lh), work in concert with sex steroids and other neurohormones to control brain function and gonadal function (Huggard et al., 1996 a, b; De Leeuw et al., 1989; Nóbrega et al., 2010; Zohar et al., 2010). In the complex and multifactorial regulation, a new hypothalamic neuropeptide, belonging to the LPXRFamide (X□=□L or Q) family, has been discovered in the brain of Japanese quail (Tsutsui et al., 2000). This peptide was named gonadotropin-inhibitory hormone (Gnih) due to its function on inhibiting the synthesis and release of gonadotropins in the pituitary of birds and mammals (Tsutsui et al., 2000; Tsutsui et al., 2010; Ubuka et al., 2012; Tsutsui et al., 2017). Unlike birds and mammals, Gnih orthologs in fish showed controversial roles in reproduction (stimulatory or inhibitory) depending on season and specie (Amano et al., 2006; Moussavi et al., 2012, 2013, 2014; Choi et al., 2016; Paullada-Salmerón et al., 2016; Muñoz-Cueto et al., 2017; Branco et., 2018; Ma et al., 2020 a, b). The stimulatory action of Gnih on gonadotropin release has been reported in sockeye salmon pituitary cells (Amano et al., 2006) and pre-spawning goldfish (Moussavi et al., 2012). However, the intraperitoneal injection of Gnih has shown to inhibit the expression of gonadotropins in the cinnamon clownfish, *Amphiprion melanopus* (Choi et al., 2016), and European sea bass, *Dicentrarchus labrax* (Paullada-Salmerón et al., 2016).

In addition to the hypothalamus, Gnih and its receptor have also been detected in the gonads of several vertebrates, including teleost fish (Bentley et al., 2008; Corchuelo et al., 2017; Muñoz-Cueto et al., 2017; Bentley et al., 2017). In zebrafish, for example, a recent study have shown that *gnih* transcripts are expressed in both germ cells and Leydig cells (Fallah et al., 2019). Moreover, it has been demonstrated that recombinant zebrafish Gnih modulated testicular function and germ cell development in zebrafish (Fallah et al., 2019,2020). Treatment with Gnih inhibited both hCG- and Fsh-induced spermatogenesis and testosterone levels in zebrafish testicular explants (Fallah et al., 2019). On the other hand, higher dose of Gnih (1000 nM) increased the basal number of haploid cells and showed paradoxical effects *in vitro* (Fallah et al., 2019). Altogether, these data support the evidence that Gnih is a player in the multifactorial regulation of reproduction, at the hypothalamus-pituitary axis by controlling gonadotropin production, as well as playing a role in the regulation of spermatogenesis and androgen production in the testes (Fallah et al., 2019, 2020; Ma et al., 2020 a, b).

In addition to the hypothalamic neuropeptides, thyroid hormones (thyroxine, T4 and triiodothyronine, T3) are also known to influence reproduction in vertebrates (Cooke et al., 1994; Buzzard et al., 2000; Maran, 2003; Swapna et al., 2006; Blanton and Specker, 2007; Habibi et al., 2012; Morais et al., 2013; Gao et al., 2014; Tovo-Neto et al., 2018; Ma et al., 2020 a, b). In mammals, thyroid hormones have been shown to regulate Sertoli cell proliferation and differentiation during testis development (Cooke et al., 1994; França et al., 1995), as well as stimulating Leydig cells differentiation and steroidogenesis in the rat testis (Mendis-Hadagama and Ariyaratne, 2005). On the other hand, information on the role of thyroid hormones in the control of male reproduction in nonmammalian vertebrates are much more limited than in mammals (Matta et al., 2002; Jacob et al., 2005; Swapna et al., 2006; Morais et al., 2013; Tovo-Neto et al., 2018; Ma et al., 2020a). In zebrafish, for example, Morais and colleagues (2013) showed that T3 increased the mitotic index of both type A undifferentiated spermatogonia (A_und_) and Sertoli cells in testicular explants. Additionally, it has been demonstrated that the stimulatory effects of T3 on the proliferation of type A_und_ spermatogonia and Sertoli cells were mediated through insulin-like growth factor signalling (Morais et al., 2013). Nonetheless, a study using the goitrogen 6-n-propyl-2-thiouracil (PTU) during the juvenile phase of Nile tilapia showed that treatment was able to increase the gonadosomatic index, Sertoli and germ cell numbers per cyst, and the number of Leydig cells the in the adult males (Matta et al., 2002). Altogether, these studies indicate the thyroid hormone regulation on germ cell proliferation is species specific and dependent on stage of development.

Although there is evidence demonstrating interaction between the brain-pituitary-gonadal and brain-pituitary-thyroid axes (Habibi et al., 2012; Tovo-Neto et al., 2018; Ma et al., 2020 a, b), much less information is available on the potential interplay of thyroid hormones with Gnih. Studies have shown that transient hypothyroidism induced by PTU in juvenile females increased *Gnih* mRNA levels and delayed onset of puberty (Kiyohara et al., 2017). Moreover, genetic loss of *Gnih* was able to reverse the PTU-induced hypothyroidism effects in female mice, demonstrating involvement of thyroid hormones on reproduction through regulation of *Gnih* expression (Kiyohara et al., 2017). The potential interaction between thyroid hormones and Gnih remains unknown in teleost fish, in particular in the gonadal function. Therefore, the aim of this study was to investigate the possible interplay of thyroid hormones with Gnih during zebrafish spermatogenesis. For this purpose, we first induced hypothyroidism in zebrafish adult males by *in vivo* treatment with methimazole, an antithyroid drug, following previously described studies (Sharma and Patiño, 2013; Sharma et al., 2016). Subsequently, *ex vivo* exposures combined with histomorphometrical and flow cytometry analysis were performed to address the possible interaction between thyroid hormones and Gnih during male germ cell development. The findings of this study provide for the first time the cellular basis for better understanding of the role of thyroid hormones in the regulation of spermatogenesis and its interaction with Gnih in the zebrafish testis.

## 2. Methods

### 2.1. Animals

Sexually mature zebrafish (4-5 months old) were bred and raised in the aquarium facility of the Department of Biological Sciences, University of Calgary, Calgary, Alberta, Canada. Fish were kept in tanks of 6-L in the recirculating system and temperature conditions similar to the natural environment (28°C) under proper photothermal conditions (14 h of light and 10 h dark). Salinity, pH, dissolved oxygen and ammonia were monitored in all tanks every day. Fish were fed twice a day using commercial food (Zeigler®). No mortality was observed during experiments. All experimental procedures were carried out according to the University of Calgary Animal Care guidelines.

### 2.2. Methimazole-induced hypothyroidism

A solution of methimazole (1-methyl-3H-imidazole-2-thione) (CAS 60-56-0; MW: 114,17; Sigma-Aldrich) was used to induce hypothyroidism. A stock solution of methimazole was prepared by dissolving 11.4 g methimazole in 1000 mL distilled water. Final concentration of methimazole (1 mM) was prepared by mixing 10 mL of the stock solution per liter of water in the tank. Two thirds of the water volume was replaced every day. The working concentration of goitrogen (1 mM) for this research was selected based on previous studies performed in our laboratory (Rodrigues, Tovo-Neto, Habibi and Nóbrega - in preparation) and by other investigators (Sharma and Patiño, 2013; Sharma et al., 2016). Zebrafish adult males were divided into control (n = 40) and methimazole treated (n = 40) groups. Fishes were exposed to methimazole for 21 days. Both groups were euthanized using buffered 2.64 g/mL MS222 tricaine mesylate (Sigma-Aldrich Oakville, Canada) at the end of treatment, and testes were dissected out for *ex vivo* organ culture experiments (see below).

### 2.3. Hormones

Zebrafish gonadotropin-inhibitory hormone (zGnih; SGTGPSATLPQRFa; MW: 1333 g/mole) was synthesized at the peptide services’ center of the University of Calgary (Calgary, Alberta, Canada). Aliquots of zGnih were reconstituted with ultrapure water and stored at -20°C. For the *in vitro* experiments, the filtered-sterilized Leibovitz’s L-15 Medium (Thermo Fisher Scientific, Gibco®, Canada) was used for dilution and preparing the final concentration of 100 nM of zGnih. Since pharmacological data are not available for zebrafish receptors, zGnih ligand concentration was chosen based on previous studies in different species including zebrafish (Moussavi et al., 2012; Branco et al., 2018; Ma et al., 2020 a, b). In most of these studies, the effects of zGnih have been seen in the range of 10-1000 nM. We preferred to not use 1000 nM zGnih because our previous study (Fallah et al., 2019) has shown that this concentration stimulated androgen production in the zebrafish testis, which in turn, triggered androgen-mediated effects on germ cells (e.g. increase of haploid cells). Human chorionic gonadotropin (hCG) as an effective Lh (Luteinizing hormone) mimicking hormone used in this study was purchased from Sigma-Aldrich (Oakville, ON, Canada). There is evidence that hCG transactivate Lhcgr (Luteinizing hormone/choriogonadotropin receptor) but not the Fshr (Follicle-stimulating hormone receptor) (Kwok et al., 2005), and *in vivo* studies in zebrafish have shown that hCG has similar effects than Lh (García-López et al., 2010). hCG was reconstituted with DNase & RNase free (ultrapure) water and mixed with filtered-sterilized L-15 to a final concentration of 5 IU/mL. Previous studies have shown that this concentration (5 IU/mL hCG) was able to stimulate zebrafish spermatogenesis increasing both testosterone levels and haploid cell (spermatids and spermatozoa) production in the testicular explants (Fallah et al., 2019, 2020).

### 2.4. Effects of zGnih in the spermatogenesis of methimazole-treated zebrafish

To investigate the effects of zGnih in the spermatogenesis of methimazole-treated zebrafish, a previously described *ex vivo* organ culture system for zebrafish testis (Leal et al., 2009) was used. In the first experiment, we evaluated the spermatogenesis progression of methimazole-treated zebrafish compared to control (non-treated fish) under basal culture conditions. For this purpose, zebrafish testes from methimazole treatment (n = 8) and control/non-treated (n = 8) were incubated with Leibovitz’s L-15 medium alone for 7 days. We then investigated the separate effects of hCG and zGnih in methimazole-treated and non-treated zebrafish spermatogenesis. Testes from methimazole treatment and control group were dissected out; one testis was incubated in Leibovitz’s L-15 Medium (basal culture medium condition) while its contra-lateral one was incubated in the presence of 5 IU/mL hCG or 100 nM zGnih for 7 days. Finally, we evaluated the effects of zGnih on hCG-stimulated spermatogenesis of zebrafish treated with the goitrogen methimazole (n = 8) and control (n = 8). For this purpose, testes were incubated with 5 IU/mL hCG in the absence or presence of 100 nM zGnih for 7 days. For all incubations, testes were placed on a nitrocellulose membrane measuring 0.25 cm^2^ (25 µm of thickness and 0.22 µm of porosity) on top of a cylinder of agarose (1.5% w/v, Ringer’s solution – pH 7.4) with 1 mL of culture medium into a 24-well plate, where medium was changed every 3 days of culture. Following the period of incubation, testes were individually collected for histomorphometrical and flow cytometry analysis (see below).

### 2.5. Histomorphometrical analysis of zebrafish spermatogenesis

Zebrafish testicular explants (n = 8 for each treatment) were fixed in modified Karnovsky (2% glutaraldehyde and 4% paraformaldehyde in Sorensen buffer [0.1 M, pH 7.2]) for at least 24 h at room temperature. Samples were subsequently dehydrated, embedded in Technovit 7100 historesin (Heraeus Kulzer, Wehrheim, Germany), sectioned at 3 µm thickness, and stained with 0,1% toluidine blue to quantify the proportion of section surface area of spermatogenic cysts containing type A undifferentiated spermatogonia (A_und_), type A differentiated (A_diff_) type B spermatogonia (SpgB), spermatocytes (Spc) and spermatids (Spt). Microscopic images of testicular tissue from five randomly chosen, non-overlapping fields of view per testicular explant were analyzed quantitatively. The analysis was based on the number of points counted over the different germ cell types investigated (Aund, Adiff, SpgB, Spc and Spt). Countings were done using a grid of 540 intersections under 100x objective lens. Germ cells were counted according to most characteristic features of each cell type i.e. nucleus structure (morphology) and cell size (Schulz et al., 2010). The proportions of section areas occupied by the different germ cell types are represented as fold change of control mean ± SEM (standard error of mean).

The relative number of spermatozoa per area was estimated as described by Fallah et al., (2019, 2020) and Tovo-Neto et al., (2020) using 20 different histological images (100x objective) per testicular explant. The images were analyzed in ImageJ software (National Institutes of Health, Bethesda, Maryland, http://rsbweb.nih.gov/ij), where background subtraction, threshold adjustment, and watershed separation of particles that resulted in a black and white picture of the highlighted spermatozoa were used to count the relative number of spermatozoa in the control and treatment groups.

### 2.6. FACscan analysis by flow cytometry (FCM)

To sort and count the number of haploid (n) cells by FACScan analysis, testicular cells from 7-days cultured tissues (n = 8 per treatment) were dissociated, fixed, and stained with red-fluorescent DNA Stain, propidium iodide (PI), as described by Fallah et al. (2019; 2020). Individual testicular explant was minced in small fragments, and testicular cells were dissociated using ethylenediaminetetraacetic acid (EDTA 5 mM) in phosphate-buffered saline (1X PBS) and Trypsin-EDTA solution 1X (Trypsin 0.05%; EDTA 0.02%). Following the dissociation, to neutralize the Trypsin-EDTA 1X solution, newborn calf serum (Sigma-Aldrich, Oakville, ON, Canada) was used. The obtained cell suspension was then centrifuged at 3500 rpm and 4°C for 15 minutes. In the following step, the cells were filtered through a 40 mm Corning cell strainer with L-15 medium to obtain a suspension of single cells. After centrifugation of the cells, they were washed in 1X PBS (Phosphate-Buffered Saline)(pH 7.4) and fixed in 70% cold ethanol for 30 minutes on a plate shaker. Afterward, to eliminate RNA for the analysis, 1X PBS supplemented with 20 mg/ml RNase A (Sigma-Aldrich, Oakville, ON, Canada) was added to the cell pellets. RNase is necessary to remove RNA-RNA or RNA-DNA double strands, which also bind to propidium iodide (PI) and can yield false positive results in FCM analysis. Cells were then stained with 12 µl of propidium iodide (PI) red-fluorescent DNA Stain from the 1 mg/ml of stock solution (Thermo Fisher Scientific, Gibco®, Canada). Following this step, to analyze the cells with a FACSCalibur system (University of Calgary equipment facilities), the prepared samples were transferred to 5 ml Polystyrene FACScan tubes. The fluorescence signals of 10000 cells from the FL2 channel of FCM were measured at 565-595 nm for each sample and blank (1X PBS supplemented with PI), and unstained cells were also used as controls. Haploid (n) cell populations (spermatids and spermatozoa) were detected by high-resolution multi-parametric flow cytometry analysis and results were analyzed using Cyflogic V1.2.1 (CyFlo Ltd., Turku, Finland), FlowJo (FlowJo LLC., Ashland, Oregon) and BD CellQuest Pro (BD Biosciences., San Jose, CA, USA). Calibration of flow cytometer was performed with the three-millimeter rainbow calibration particles (Spherotech, Libertyville, IL) before each run. The haploid cell number (spermatids and spermatozoa) for each treatment was represented as percentage of fold to control (basal culture medium condition).

### 2.7. Statistical analysis

Values are presented as mean ± SEM, and data were analyzed using paired t-test since they represent paired observations for each individual (one testis served as control to the other one treated). However, when comparing three or more groups of testis explants from FCM results were evaluated by one-way ANOVA followed by Tukey’s post-hoc multiple comparisons test. The GraphPad Prism V6.0 (Graphpad Software Inc., La Jolla, CA, USA) was used for all of the statistical analyses. Significance level (p) was fixed at 0.05 in all experiments.

## 3. Results

### Effects of methimazole on basal spermatogenesis by ex vivo approach

After *in vivo* treatment with the goitrogen methimazole (1mM for 21 days exposure), zebrafish testes were incubated with L-15 for 7 days to assess progression of spermatogenesis under basal culture conditions (Fig. 1B). Testes from untreated fish (control) was incubated in parallel as control for 7 days (Fig. 1A). Histomorphometrical analysis revealed that methimazole treatment affected the proportion of different germ cell cysts after 7 days of culture, compared to control (Fig. 1C). In this context, there is a significant increase in the proportion of the area occupied by type A differentiated spermatogonia (A_diff_), type B spermatogonia and spermatocytes in the testis of fish treated with methimazole, compared to untreated fish (Fig. 1C). In the methimazole treated fish, however, there were reduction in the proportion area occupied by spermatids (Fig. 1C), basal number of spermatozoa (Figs. 2A-B, 3D), and haploid cell population (spermatids and spermatozoa) (Fig. 4), compared to control as determined by histomorphometrical and FCM analysis.

**Figure 1.**
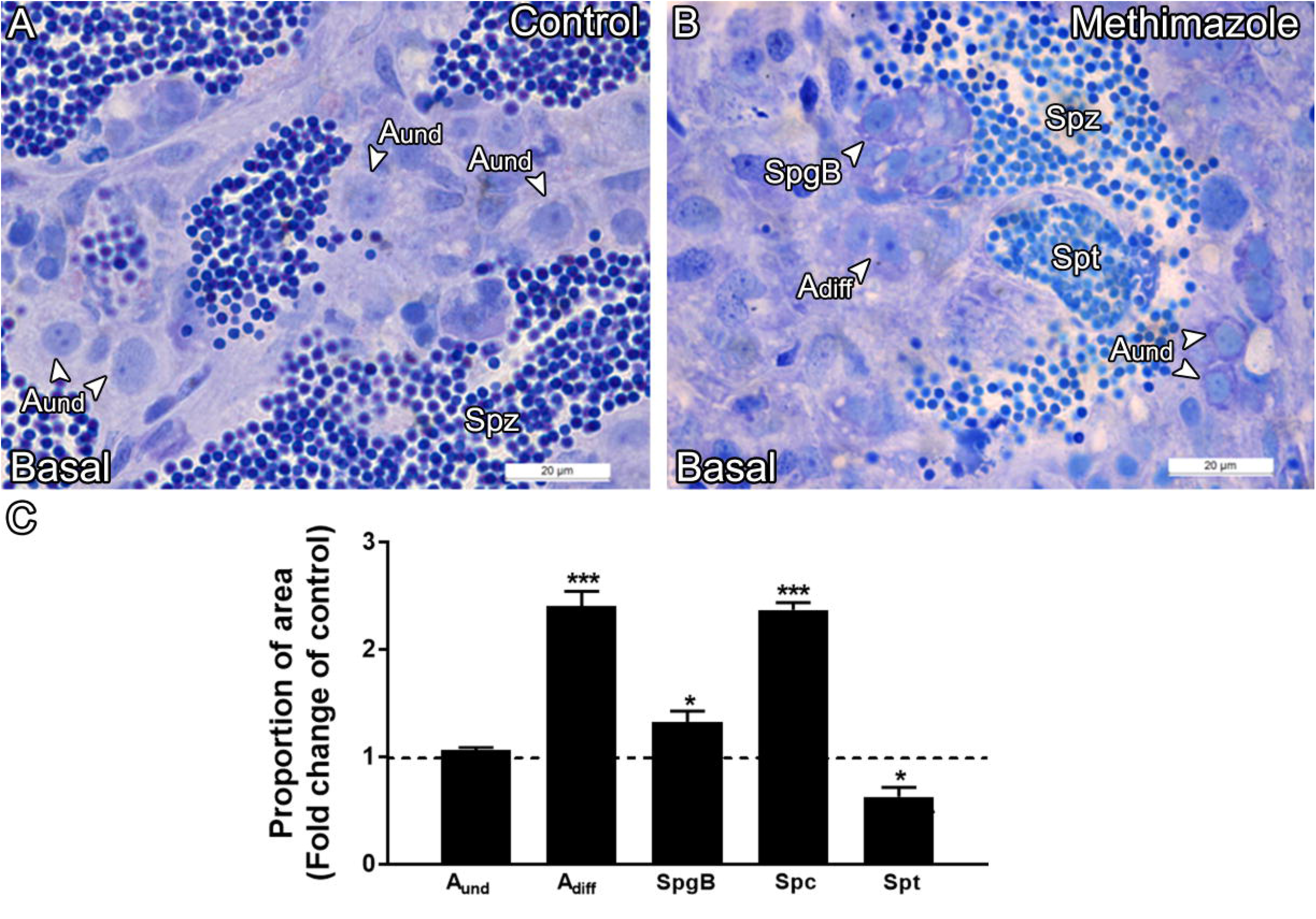
Histomorphometrical analysis of zebrafish testis incubated for 7 days with L-15 medium (basal culture medium) following *in vivo* treatment with the goitrogen methimazole (1mM for 21 days exposure). (**A):** Control group (non-treated fish) (n = 8). **(B):** Methimazole-treated group (n = 8). Type A undifferentiated spermatogonia (A_und_), type A differentiated spermatogonia (A_diff_), type B spermatogonia (SpgB), spermatocytes (Spc) and spermatids (Spt). Staining: Toluidine blue. Scale bar = 20 µm. **(C):** Proportion of section area occupied by spermatogenic cysts containing A_und_, A_diff_, SpgB, Spc or Spt in testicular explants from methimazole treated group incubated for 7 days with L-15 medium (basal culture medium). Values (mean ± SEM) are expressed as fold change of the untreated group (control) (dotted line set at 1). Asterisks denote statistical significance between the control and methimazole group; *\* p < 0*.*05*; *\*\*\* p < 0*.*001* (Student unpaired *t*-test; n = 8).

**Figure 2.**
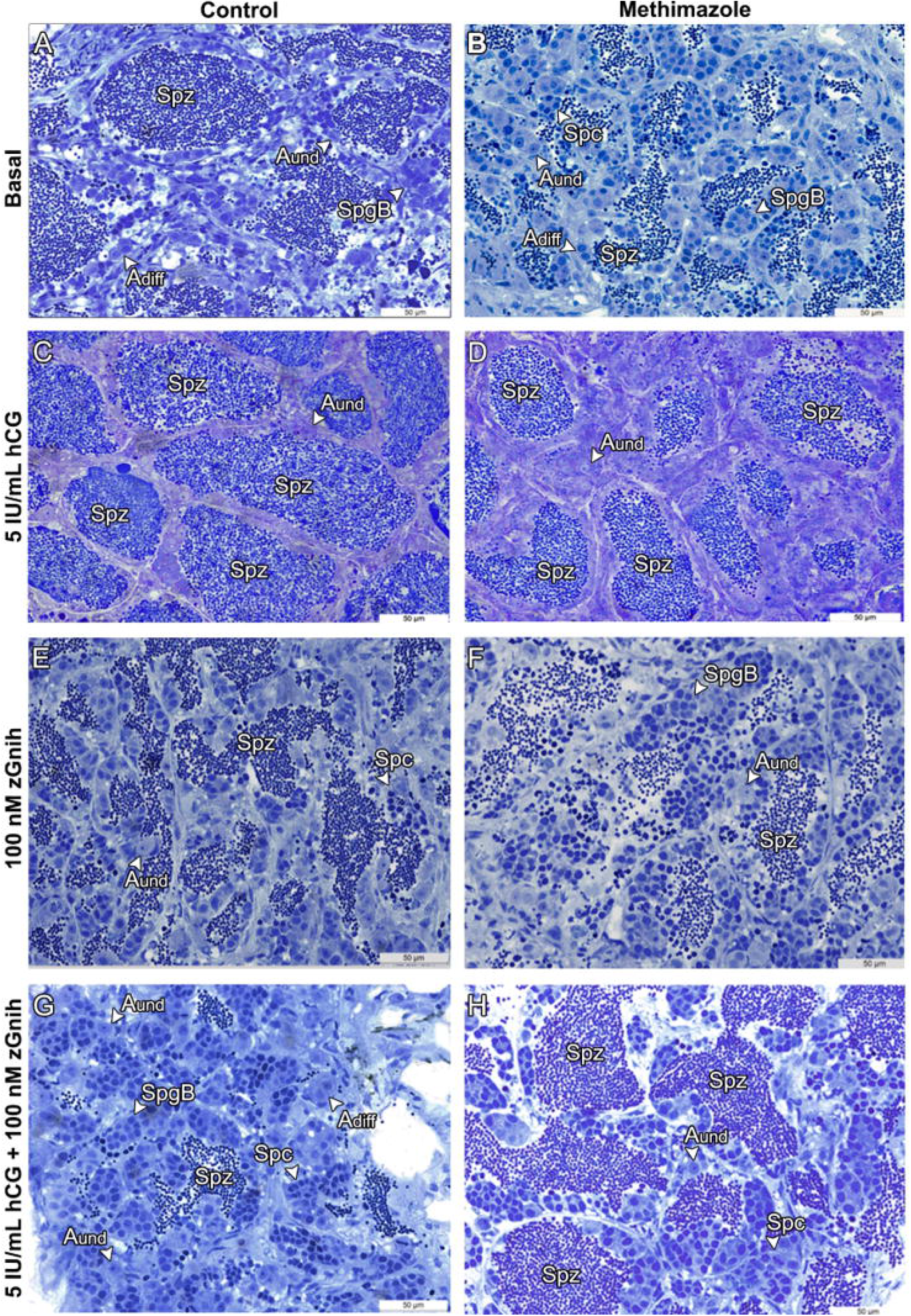
Morphological analysis of zebrafish testicular explants from untreated group (left column) and methimazole-treatment (right column), after 7 days in culture, under different conditions. **(A-B):** basal culture medium (L-15); **(C-D):** 5 IU/mL hCG; **(E-F):** 100 nM zGnih; **(G-H):** 5 IU/mL hCG + 100 nM zGnih. Type A undifferentiated spermatogonia (A_und_), type A differentiated spermatogonia (A_diff_), type B spermatogonia (SpgB), spermatocytes (Spc), spermatids (Spt) and spermatozoa (Spz) are indicated. Staining: Toluidine blue. Scale bar = 50 µm.

### Effects of methimazole on hCG-induced spermatogenesis

We also investigated the effects of methimazole treatment (1mM for 21 days exposure) on hCG-induced spermatogenesis by incubating testes from treated and control zebrafish for 7 days in L-15 medium (Fig. 2A-D). Histomorphometrical analysis revealed that hCG (5 IU/mL) did not change the basal proportion of early, meiotic or post-meiotic germ cells (types A_und_, A_diff_ and B spermatogonia, spermatocytes and spermatids) (Fig. 3A). However, methimazole treatment significantly decreased the proportion area occupied by type B spermatogonia, and increased the proportion area occupied by spermatids in the zebrafish testis, compared to control (Fig. 3A). As expected, hCG (5 IU/mL) increased the basal number of spermatozoa (Figs. 2A-D and 3D) and haploid cell population (spermatids and spermatozoa) (Fig. 4) in both groups (control and methimazole). However, the potency of hCG in terms of inducing haploid cell production was reduced in the testis of zebrafish treated with methimazole, compared to control (Figs. 2A-D, 3D, 4).

**Figure 3.**
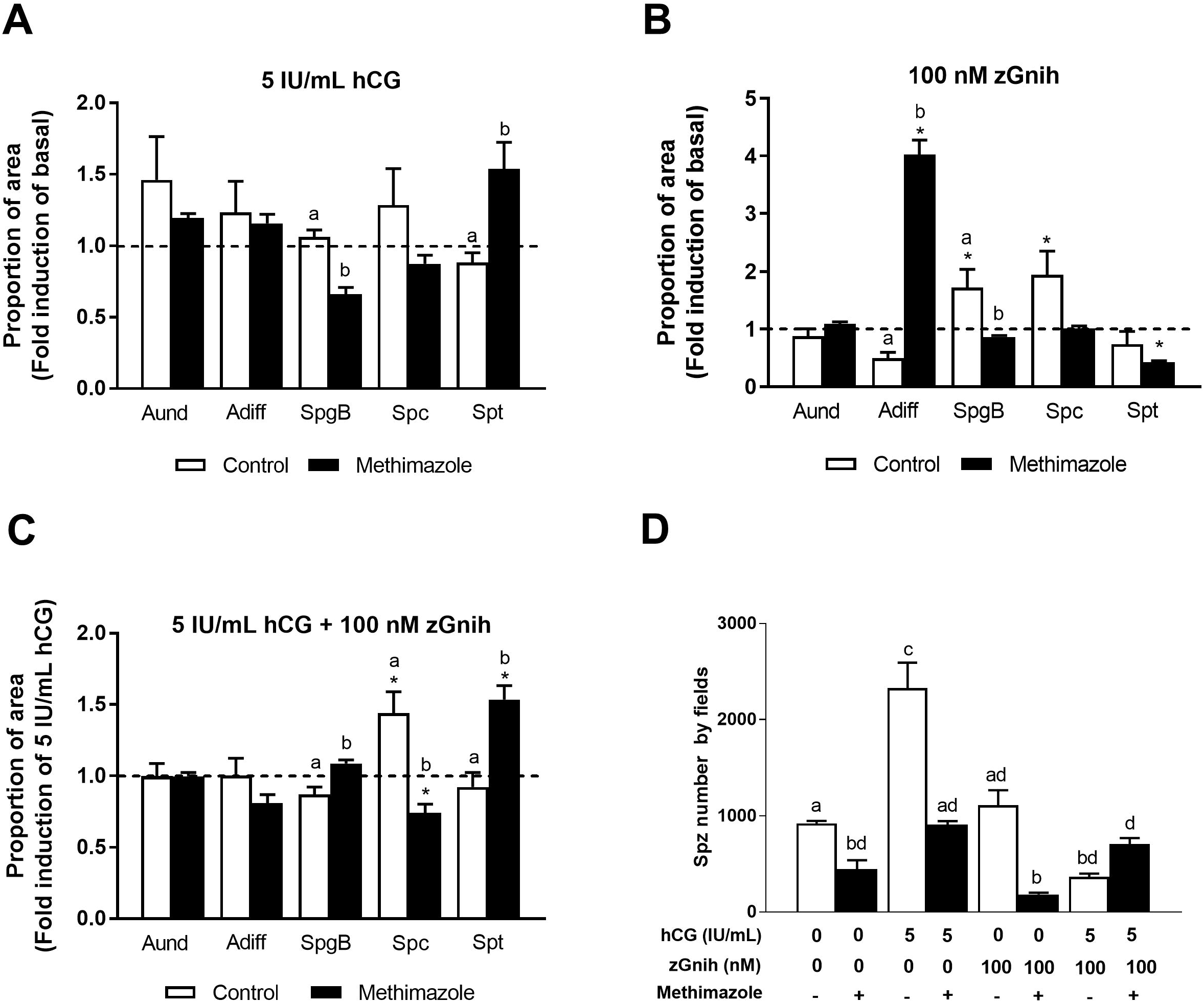
Effects of hCG (5 IU/mL), zGnih (100 nM), or hCG (5 IU/mL) + zGnih (100 nM) on zebrafish spermatogenesis from control (non-treated) and methimazole treatment after 7 days in culture. **(A-C):** Proportion of section surface area occupied by cysts of type A undifferentiated spermatogonia (A_und_), type A differentiated (A_diff_), type B spermatogonia (SpgB), spermatocytes (Spc) and spermatids (Spt) in the testicular explants from control/untreated fish (white bars) and methimazole (black bars) treated with hCG (A), zGnih (B) or hCG in the presence of zGnih (C). Values (mean ± SEM) (n = 8 testes per condition) are presented as fold change of basal control condition: L-15/basal medium condition **(A**,**B)** or hCG incubation **(C)**, set at 1 (dotted line). Asterisks denote statistical significance from basal control condition [L-15/basal medium condition **(A**,**B)** or hCG incubation **(C)**]; *\* p < 0*.*05* (Student’s paired *t*-test). Different letters denote significant differences among groups (non-treated and methimazole treatment); *p < 0*.*05* (ANOVA followed by Tukey’s multiple comparison test). **(D)** Spermatozoa number by fields from control (white bars) and methimazole (black bars) testicular explants after 7 days in culture under different conditions. Bars represent mean ± SEM (ANOVA followed by Tukey’s multiple comparison test; n = 8). Different letters denote statistically differences (p < 0.05) between different treatment conditions.

**Figure 4.**
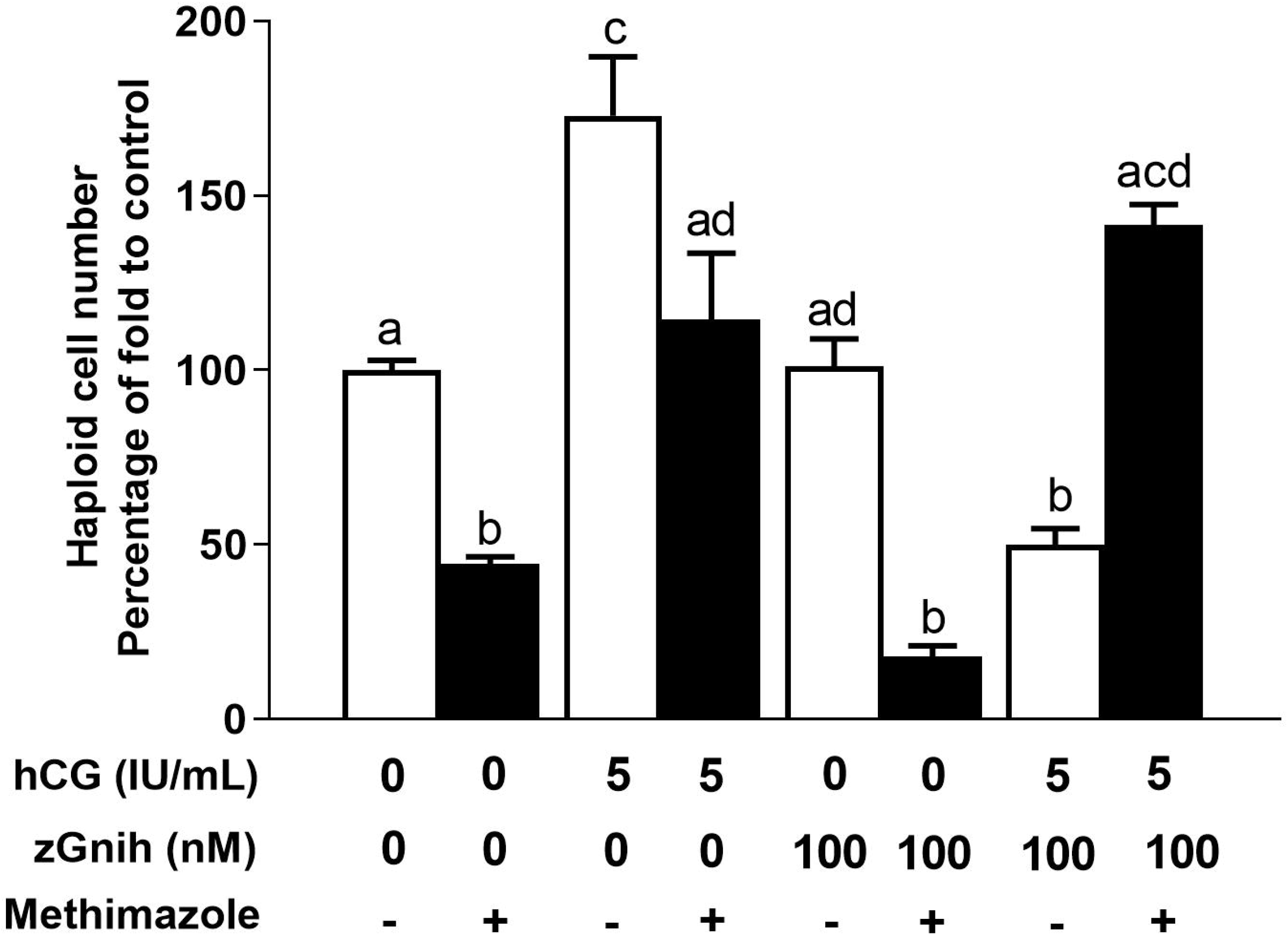
Effects of hCG (5 IU/mL), zGnih (100 nM) or hCG (5 IU/mL) + zGnih (100 nM) on zebrafish testicular haploid cell population (spermatids and spermatozoa) from control (non-treated) and methimazole treatment after 7 days in culture. Following 7 days in culture under different conditions (n = 8 testes per condition), zebrafish testicular explants from control (white bars) and methimazole treatment (black bars) were dissociated and assayed for cell cycle analysis. The percentage of haploid cell was determined by FACScan analysis. Counts were normalized to 100%; treatment groups were normalized against control (0 hcG + 0 zGnih) and expressed as mean ± SEM. Different letters denote significant differences among groups; *p < 0*.*05* (ANOVA followed by Tukey’s multiple comparison test).

### Effects of methimazole on zGnih mediated actions on basal and hCG-induced spermatogenesis

To investigate the effects of methimazole on zGnih mediated zebrafish spermatogenesis, testes of the goitrogen methimazole (1mM for 21 days exposure) and untreated control were incubated for 7 days in L-15 in the following conditions: 1) presence and absence of zGnih (100 nM), and 2) presence of hCG (5 IU/mL) with and without zGnih (100 nM) (Fig. 2). In these studies, methimazole treatment significantly altered zGnih-induced germ cell proportion as compared to control (Fig. 3B). Histomorphometrical analysis revealed that treatment with methimazole significantly increased zGnIH induced proportion area occupied by type A_diff_, and decreased the area occupied by type B spermatogonia with respect to untreated control fish (Fig. 3B). Although, histomorphometrical and FCM analysis revealed lower number of spermatozoa (Figs. 2E,F, 3D) and haploid cell population (Fig. 4) following methimazole treatment, zGnih did not modulate the basal spermatozoa production in both experimental groups (methimazole and control)(Fig. 3D, 4). Subsequently, we investigated the effects of methimazole treatment on zGnih mediated actions on hCG-induced spermatogenesis. Previous studies have shown that zGnih inhibits the hCG-induced spermatogenesis in zebrafish testis (Fallah et al., 2019). The present study demonstrated that methimazole significantly alter the inhibitory action of zGnih on hCG-induced spermatogenesis in the zebrafish testis (Figs. 2G,H, 3C,D, 4). In the methimazole treated fish, zGnih in the presence of hCG significantly increased the proportion area of type B spermatogonia and spermatids, compared to control groups (Fig. 3C). Conversely, combined treatments with zGnih and hCG decreased the proportion area occupied by spermatocytes in the testis of zebrafish treated with methimazole (Fig. 3C). Furthermore, histomorphometrical and FCM analysis revealed that methimazole treatment nullify the inhibitory action of zGnih on the hCG-induced spermatozoa and haploid cell population (Figs. 3D, 4).

## 4. Discussion

Reproduction is under multifactorial control of neurohormones, pituitary hormones, glucocorticoids, growth factors, thyroid hormones as well as a number of gonadal hormones including steroids and peptides (Schulz et al., 2010; Tovo-Neto et al., 2018; Fallah 2019, 2020, 2021; Ma et al., 2020a, b; Tovo-Neto et al., 2020; Li et al., 2020). A number of neurohormones including Gnih (Tsutsui et al., 2000) were shown to be involved in the control of pituitary gonadotropin production (Amano et al., 2006; Moussavi et al., 2012, 2013, 2014; Choi et al., 2016; Paullada-Salmerón et al., 2016; Muñoz-Cueto et al., 2017; Branco et., 2018; Ma et al., 2020 a, b) as well as being involved in paracrine control of gonadal function in fish (Fallah et al., 2019, 2020, 2021). A number of studies have shown that thyroid hormones interact with the reproductive axis (Habibi et al., 2012) and are, in particular, involved in the regulation of the testicular function in fish (Matta et al., 2002; Jacob et al., 2005; Swapna et al., 2006; Morais et al., 2013; Tovo-Neto et al., 2018). Based on this background, we investigated the interaction between Gnih and thyroid hormones in the control of spermatogenesis in adult zebrafish. To this end, we exposed zebrafish males to the goitrogen methimazole (1mM for 21 days), which is known to induce hypothyroidism by blocking thyroid peroxidase (TPO) and thyroid hormone synthesis (Sharma and Patiño, 2013; Sharma et al., 2016; Rodrigues, Tovo-Neto, Habibi and Nóbrega - in preparation). Subsequently, a factorial design using an *ex vivo* organ culture system for zebrafish testis (Leal et al., 2009) was adopted to investigate the interaction between Gnih and thyroid hormones during germ cell development. In this study, we first evaluated the effects of methimazole on basal spermatogenesis using histomorphometric and FCM analysis. These analyses showed that methimazole treatment affected zebrafish germ cell development by increasing the proportion area occupied by type A differentiated spermatogonia (A_diff_), type B spermatogonia and spermatocytes, while spermatids and spermatozoa were significantly decreased, compared to control. Our results demonstrate that thyroid hormones are required for normal germ cell development for both meiotic and post-meiotic cells, as shown by the increase in spermatocytes and decrease in the haploid cells (spermatids and spermatozoa), respectively. Our results are compatible with an earlier study demonstrating that treatment with thiourea, another goitrogen, decrease the number of spermatids and spermatozoa in the adult testes of African catfish (*Clarias gariepinus*) (Swapna et al., 2006). Our results are also compatible with the reports that hypothyroidism can decrease fertility in men (Krajewska-Kulak and Sengupta 2013). Although the mechanisms underlying the reduced fertility are unclear, it is possible that thyroid hormone deficiency may have affected steroidogenesis and androgen sensitivity, which, consequently, had an impact in the activity of the hypothalamic-pituitary-gonadal axis. Also in agreement with our result a previous study demonstrated that treatment with thyroid hormones potentiate gonadotropin-induced androgen production and increase the expression of androgen receptor in the zebrafish testis (Morais et al., 2013).

Our results demonstrate increases in the proportion area of all types of spermatogonia cells after treatment with methimazole, except for type A undifferentiated (A_und_) spermatogonia which remained unaffected, compared to control. This is consistent with the observation that methimazole-induced hypothyroidism significantly impair meiosis and spermiogenesis, leading to accumulation of pre-meiotic cysts in the zebrafish explants. In this context, a previous study demonstrated that T3 (50 ng/mL) increased the proportion area and mitotic index of type A_und_ spermatogonia in the zebrafish testicular explants (Morais et al., 2013). Furthermore, it was shown that Igf3 (Insulin-like growth factor 3) which is produced by Sertoli cells (Nóbrega et al., 2010), mediate T3-induced proliferation of type A_und_ spermatogonia (Morais et al., 2013). Therefore, as expected a lower proliferation rate of type A_und_ spermatogonia was observed following methimazole-induced hypothyroidism in the present study. The low mitotic index, in combination with the meiosis impairment, could explain the unaltered proportion area occupied for type A_und_ spermatogonia when compared to control. Overall, the results of present study support the hypothesis that thyroid hormones are important for normal testicular development and spermatogenesis and hypothyroidism can adversely affect germ cell development in adult zebrafish. Species specificity and complex nature of thyroid hormone on testicular development was also demonstrated following induction of transitory hypothyroidism in Nile tilapia (*Oreochromis niloticus*), leading to elevated gonadosomatic index as well as increased number of Sertoli and germ cells per cyst in the testis (Matta et al., 2002). While there is a strong evidence for crosstalk between thyroid hormones and different components of reproductive axis, there are variation in thyroid response depending on species and stage of development.

In the next step, we investigated potential interaction between thyroid hormones and gonadotropin-induced response. We used hCG as an effective Lh (Luteinizing hormone) mimicking hormone since both hormones have similar effects in the zebrafish testes (García-López et al., 2010). As expected, treatment with hCG (5 IU/ml) increased the number of haploid cells, in particular of spermatozoa, in the zebrafish testes (positive control). This result is in line with a recent study demonstrating stimulatory effect of hCG (5 IU/mL) on the spermatozoa production and testosterone release in the zebrafish testes (Fallah et al., 2019). However, treatment with methimazole significantly altered the hCG-induced haploid cell numbers, indicating that thyroid hormones are essential for normal gonadotropin function in the zebrafish testes. Similar results describing the interaction between thyroid hormones and hCG/LH has been reported in mammals before. For example, methimazole-induced hypothyroidism in immature rats reduced the hCG/LH bindings sites in the testis, affected Leydig cell responsiveness to hCG stimulation, and consequently, impaired testosterone production *in vitro* (Valle et al., 1985). Similarly, a study with cultured human luteinized granulosa cells has demonstrated that thyroid hormones modulate the hCG-mediated functions in ovarian cells, highlighting the influence of thyroid hormones on gonadotropin action (Goldman et al., 1993). In adult testes, Manna and collaborators (2001) showed that thyroid hormones influence the expression of LH/hCG receptors using hypo- and hyperthyroid mice. Altogether, these results are consistent with the present observation that methimazole-induced hypothyroidism impair gonadotropin-induced response in the zebrafish testes. Likewise, it is possible that thyroid hormones may also regulate Lh/choriogonadotropin receptor (Lhcgr) expression in zebrafish testes. In the zebrafish testes, Lhcgr is expressed in Leydig (García-López et al., 2010) and germ cells (spermatocytes and round spermatids) (Chauvigné et al., 2014). Moreover, it has been shown in zebrafish that Lhcgr activation in Leydig cells results in androgen release (García-López et al., 2010), whereas in germ cells, this receptor mediate more direct action of Lh/hCG on spermiogenesis (Chauvigné et al., 2014). Therefore, deficiency of thyroid hormone affects both Leydig cell function (steroidogenesis), and Lh/hCG-induced spermiogenesis in zebrafish testes. In this context, histomorphometrical analysis revealed that methimazole treatment reduce hCG-induced increase in the proportion area of spermatids leading to reduced number of spermatozoa. Nevertheless, more studies will be necessary to elucidate the molecular basis underlying the interaction between thyroid hormones and Lh/hCG in the zebrafish testes.

After investigating the effects of methimazole on basal and hCG-induced spermatogenesis, we investigated the interplay between thyroid hormones and Gnih through the same approach using the *ex vivo* system. Gnih was discovered in the brain of Japanese quail as a new member of the LPXRFamide family that inhibited the gonadotropin release in the pituitary (Tsutsui et al., 2000; Tsutsui et al., 2010; Ubuka et al., 2012; Tsutsui et al., 2017). Among teleosts, Gnih orthologs have been identified in a number of species, and was shown to have both stimulatory and inhibitory actions, depending on season (Amano et al., 2006; Moussavi et al., 2012, 2013, 2014; Choi et al., 2016; Paullada-Salmerón et al., 2016; Muñoz-Cueto et al., 2017; Branco et., 2018; Ma et al., 2020 a, b). The presence of Gnih and its receptors have been shown in the gonads of vertebrates (Bentley et al., 2008; Corchuelo et al., 2017; Muñoz-Cueto et al., 2017; Bentley et al., 2017), providing supporting evidence for a local, paracrine/autocrine role of Gnih in the gonads. More recent studies in zebrafish demonstrated that *gnih* is widely expressed among the testicular germ cells, and also in the interstitial compartment, especially in the Leydig cells (Fallah et al., 2019). Moreover, it was shown that zebrafish Gnih (100 nM) inhibited both hCG-, and Fsh-induced androgen release and haploid cell production (spermatids and spermatozoa) in the zebrafish testis (Fallah et al., 2019, 2020). The present study demonstrates for the first time that Gnih and thyroid hormones are part of an orchestrated multifactorial mechanism that regulate testicular function and germ cell development in zebrafish. Our results demonstrate that methimazole-induced hypothyroidism effect the Gnih-mediated actions on basal and hCG-induced spermatogenesis in the zebrafish testis. We observed that methimazole-induced hypothyroidism nullify the basal effects of Gnih in the spermatogonial phase, more specifically reducing the proportion area of type B spermatogonia and increasing the area occupied by type A differentiated (A_diff_) spermatogonia, compared to control. We postulate that the observed results were likely due to decreased proliferation rate of type B spermatogonia, which led to increased accumulation of type A_diff_ spermatogonia in the testes. With respect to the haploid cell population, although methimazole had lower number of haploid cells (spermatids and spermatozoa) when compared to control, Gnih did not modulate the basal production of these cells in both groups (methimazole and untreated fish). These results are in agreement with Fallah and colleagues (2019) who demonstrated that treatment with zebrafish Gnih (100 nM) did not affect the basal number of spermatozoa and testosterone. With regard to hCG-induced spermatogenesis, goitrogen treatment nullified the inhibitory effects of Gnih on the hCG-induced spermatozoa, and altered the proportion area occupied by germ cells following co-treatment with Gnih and hCG. We also observed an increase in type B spermatogonia, reduction of spermatocytes, and augmentation of spermatids in the methimazole-treated group, compared to control.

Overall, our data clearly demonstrate that thyroid hormones are necessary for Gnih mediated functions on basal and hCG-induced spermatogenesis in the zebrafish testis. The most prominent effect was on gonadotropin-induced spermatogenesis, in which the inhibitory action of Gnih on hCG-induced testicular response was blocked in methimazole-induced hypothyroid zebrafish. It should be noted that the observed local effects on testicular development is compounded by central effects of thyroid hormones on the reproductive axis (Habibi et al., 2012). In the brain, Kiyohara and colleagues (2017) demonstrated that hypothyroidism induced by PTU in juvenile female mice leads to delayed pubertal onset by increasing *Gnih* expression leading to reduced pituitary gonadotropin (LH) and E2 levels. Therefore, further studies will be necessary to elucidate the integrated effects of thyroid hormones on Gnih mediated functions in zebrafish brain-pituitary-gonadal axis.

In summary, our findings provide novel information on the control of zebrafish spermatogenesis and interaction between the brain-pituitary-gonadal and brain-pituitary-thyroid axes. We have demonstrated that thyroid hormones interaction with hCG and Gnih are important components of multifactorial regulation of spermatogenesis in the zebrafish testis, and absence of thyroid hormones leads to impairment of testicular function and germ cell development in zebrafish. The findings from the present study, provides a framework for better understanding of the role of thyroid hormones in the control of reproduction and testicular function in fish and other vertebrates.

## Author contributions

R.H.N., M.S.R., and H.R.H. designed the study; M.S.R., H.P.F., and M.Z. performed the experiments; All authors analyzed the data; R.H.N., M.S.R. and H.R.H. wrote the paper. All authors edited the article.

## References

Amano, M., Moriyama, S., Ligo, M., Kitamura, S., Amiya, N., Yamamori, K., Ukena, K., Tsutsui, K., 2006. Novel fish hypothalamic neuropeptides stimulate the release of gonadotrophins and growth hormone from the pituitary of sockeye salmon. J. Endocrinol. 188, 417–423. https://doi.org/10.1677/joe.1.06494.

Bentley, G.E., Ubuka, T., McGuire, N.L., Chowdhury, V.S., Morita, Y., Yano, T., Hasunuma, I., Binns, M., Wingfield, J.C., Tsutsui, K., 2008. Gonadotropin-inhibitory hormone and its receptor in the avian reproductive system. Gen Comp Endocrinol. 156, 34–43. https://doi.org/10.1016/j.ygcen.2007.10.003.

Bentley, G.E., Wilsterman, K., Ernst, D.K., Lynn, S.E., Dickens, M.J., Calisi, R.M., Kriegsfeld, L.J., Kaufer, D., Geraghty, A.C., viviD, D., McGuire, N.L., Lopes, P.C., Tsutsui K., 2017. Neural Versus Gonadal GnIH: Are they Independent Systems? A Mini-Review. Integr Comp Biol. 1;57(6):1194–1203. https://doi.org/10.1093/icb/icx085.

Branco, G.S., Melo, A.G., Ricci, J.M.B., Digmayer, M., de Jesus, L.W.O., Habibi, H.R., Nóbrega, R.H., 2018. Effects of GnRH and the dual regulatory actions of GnIH in the pituitary explants and brain slices of Astyanax altiparanae males. Gen. Comp. Endocrinol. 273, 209–217. https://doi.org/10.1016/j.ygcen.2018.08.006.

Blanton, M.L., Specker, J.L., 2007. The hypothalamic-pituitary-thyroid (HPT) axis in fish and its role in fish development and reproduction. Crit. Rev. Toxicol. 37, 97–115. https://doi.org/10.1080/10408440601123529.

Buzzard, J.J., Morrison, J.R., O’Bryan, M.K., Song, Q., Wreford, N.G., 2000. Developmental expression of thyroid hormone receptors in the rat testis. Biol. Reprod. 62, 664–669. https://doi.org/10.1095/biolreprod62.3.664.

Chauvigné, F., Zapater, C., Gasol, J.M., Cerda, J., 2014. Germ-line activation of the luteinizing hormone receptor directly drives spermiogenesis in a nonmammalian vertebrate. Proceedings of the National Academy of Sciences. 111, 1427–1432. https://doi.org/10.1073/pnas.1317838111.

Choi, Y.J., Kim, N.N., Habibi, H.R., Choi, C.Y., 2016. Effects of gonadotropin inhibitory hormone or gonadotropin-releasing hormone on reproduction-related genes in the protandrous cinnamon clownfish, Amphiprion melanopus. Gen. Comp. Endocrinol. 235, 89–99. https://doi.org/10.1016/j.ygcen.2016.06.010.

Cooke, P.S., Zhao, Y.D., Bunick, D., 1994. Triiodothyronine inhibits proliferation and stimulates differentiation of cultured neonatal Sertoli cells: possible mechanism for increased adult testis weight and sperm production induced by neonatal goitrogen treatment. Biol. Reprod. 51, 1000–1005. https://doi.org/10.1095/biolreprod51.5.1000.

Corchuelo, S., Martinez, E.R.M., Butzge, A.J., Doretto, L.B., Ricci, J.M.B., Valentin, F.N., Nakaghi, L.S.O., Somoza, G.M., Nóbrega, R.H., 2017. Characterization of Gnrh/Gnih elements in the olfacto-retinal system and ovary during zebrafish ovarian maturation. Mol Cell Endocrinol. 450, 1–13. https://doi.org/10.1016/j.mce.2017.04.002.

De Leeuw, R., Habibi, H.R., Nahorniak, C.S., Peter, R.E., 1989. Dopaminergic regulation of pituitary gonadotrophin-releasing hormone receptor activity in the goldfish (Carassius auratus). J. Endocrinol. 121, 239–247. https://doi.org/10.1677/joe.0.1210239.

Fallah, H.P., Tovo-Neto, A., Yeung, E.C., Nóbrega, R.H., Habibi, H.R., 2019. Paracrine/ autocrine control of spermatogenesis by gonadotropin-inhibitory hormone. Mol. Cell. Endocrinol. 492, 110440. https://doi.org/10.1016/j.mce.2019.04.020.

Fallah, H. P., Rodrigues, M. S., Corchuelo, S., Nóbrega, R. H., Habibi, H. R., 2020. Role of GnRH isoforms in paracrine/autocrine control of zebrafish (Danio rerio) spermatogenesis. Endocrinol. https://doi.org/10.1210/endocr/bqaa004.

Fallah, H.P., Rodrigues, M.S., Zanardini, M., Nóbrega, R.H., Habibi, H.R., 2021. Effects of gonadotropin-inhibitory hormone on early and late stages of spermatogenesis in ex-vivo culture of zebrafish testis. Mol Cell Endocrinol. 15;520:111087. https://doi.org/10.1016/j.mce.2020.111087.

França, L.R., Hess, C.A., Cooke, P.S., Russel, L.D., 1995. Neonatal hypothyroidism causes delayed Sertoli cell maturation in rats treated with propylthiouracil: evidence that the Sertoli cell controls testis growth. Anat. Rec. 242, 57–69. https://doi.org/10.1002/ar.1092420108.

Gao, Y., Lee, W. M., Cheng, C. Y., 2014. Thyroid hormone function in the rat testis. Front. Endocrinol. 5:188. https://doi.org/10.3389/fendo.2014.00188.

García-López, A., Jonge, H., Nóbrega, R.H., Waal, P.P., Van Dijk, W., Hemrika, W., Taranger, G.L., Bogerd, J., Schulz, R.W., 2010. Studies in zebrafish reveal unusual cellular expression patterns of gonadotropin receptor messenger ribonucleic acids in the testis and unexpected functional differentiation of the gonadotropins. Endocrinol. 151, 2349–2360. https://doi.org/10.1210/en.2009-1227.

Goldman, S., Dirnfeld, M., Abramovici, H., Kraiem, Z., 1993. Triiodothyronine (T3) modulates hCG-regulated progesterone secretion, cAMP accumulation and DNA content in cultured human luteinized granulosa cells. 125–131. https://doi.org/10.1016/0303-7207(93)90102-P.

Habibi, H.R., Andreu-Vieyra, C.V., 2007. Hormonal regulation of follicular atresia in teleost fish. The fish Oocyte: From basic Studies to Biotechnological Applications. 235–253. https://doi.org/10.1007/978-1-4020-6235-3_9.

Habibi, H.R., Nelson, E.R., Allan, E.R.O., 2012. New insights into thyroid hormone function and modulation of reproduction in goldfish. Gen. Comp. Endocrinol. 175, 19–26. https://doi.org/10.1016/j.ygcen.2011.11.003.

Huggard, D., Khakoo, Z., Kassam, G., Seyed Mahmoud, S., Habibi, H. R., 1996a. Effect of Testosterone on Maturational Gonadotropin Subunit Messenger Ribonucleic Acid Levels in the Goldfish Pituitary. Biol. Reprod. 54, 1184–1191. https://doi.org/10.1095/biolreprod54.6.1184.

Huggard, D., Khakoo, Z., Kassam, G., Habibi, H. R., 1996b. Effect of testosterone on growth hormone gene expression in the goldfish pituitary. Can. J. Physiol. Pharmacol. 74, 1039–1046. https://doi.org/10.1139/y96-103.

Jacob, T.N., Pandey, J.P., Raghuveer, K., 2005. Thyroxine-induced alterations in the testis and seminal vesicles of air-breathing catfish, Clarias gariepinus. Fish Physiol. Biochem. 31, 271. https://doi.org/10.1007/s10695-006-0035-0.

Kiyohara, M., Son, Y.L., Tsutsui, K., 2017. Involvement of gonadotropin-inhibitory hormone in pubertal disorders induced by thyroid status. Sci. Rep. 7, 1042. https://doi.org/10.1038/s41598-017-01183-8.

Krajewska-Kulak, E., Sengupta, P., 2013. Thyroid function in male infertility. Front. Endocrinol. 4, 174. https://doi.org/10.3389/fendo.2013.00174.

Kwok, H.-F., So, W.-K., Wang, Y., Ge, W., 2005. Zebrafish gonadotropins and their receptors: I. cloning and characterization of zebrafish follicle-stimulating hormone and luteinizing hormone receptors-evidence for their distinct functions in follicle development. Biol. Reprod. 72, 1370–1381. https://doi.org/10.1095/biolreprod.104.038190.

Li, M., Liu, X., Dai, S., Xiao, H., Qi, S., Li, Y., Zheng, Q., Jie, M., Cheng, C.H.K., Wang, D., 2020. Regulation of spermatogenesis and reproductive capacity by Igf3 in tilapia. Cell. Mol. Life Sci. 77, 4921–4938. https://doi.org/10.1007/s00018-019-03439-0.

Leal, M.C., De WaaL, P.P., García-López, A., Chen, S.X., Bogerd, J., Schulz, R.W., 2009. Zebrafish primary testis tissue culture: An approach to study testis function ex vivo. Gen. Com. Endocrinol. 162, 134–138. https://doi.org/10.1016/j.ygcen.2009.03.003.

Ma, Y., Ladisa, C., Chang, J.P., Habibi, H. R., 2020a. Multifactorial control of reproductive and growth axis in male goldfish: Influences of GnRH, GnIH and thyroid hormone. Mol. Cell. Endocrinol. 500, 110629. https://doi.org/10.1016/j.mce.2019.110629.

Ma, Y., Ladisa, C., Chang, J.P., Habibi, H.R., 2020b. Seasonal Related Multifactorial Control of Pituitary Gonadotropin and Growth Hormone in Female Goldfish: Influences of Neuropeptides and Thyroid Hormone. Front. Endocrinol. 11. https://doi.org/10.3389/fendo.2020.00175.

Manna, P.R., Roy, P., Clark, B.J., Stocco, D.M., Huhtaniemi, I.T., 2001. Interaction of thyroid hormone and steroidogenic acute regulatory (StAR) protein in the regulation of murine Leydig cell steroidogenesis. J. Steroid. Bioch. Mol. Biol. 76, 167–177. https://doi.org/10.1016/S0960-0760(00)00156-4.

Maran, R.R.M., 2003. Thyroid hormones: their role in testicular steroidogenesis, Arch. Androl. 49, 375–388. https://doi.org/10.1080/01485010390204968.

Matta, S.L, Vilela, D.A, Godinho, H.P, França, L.R., 2002. The goitrogen 6-n-propyl-2-thiouracil (PTU) given during testis development increases Sertoli and germ cell numbers per cyst in fish: the tilapia (Oreochromis niloticus) model. Endocrinol. 143, 970–978. https://doi.org/10.1210/endo.143.3.8666.

Mendis-Handagama, S.M.L.C., Ariyaratne, H.B.S., 2005. Leydig cells, thyroid hormones and steroidogenesis. Indian J. Exper. Biol. 43, 939–962.

Morais, R.D., Nóbrega, R.H., Gómez-González, N.E., Schmidt, R., Bogerd, J., França, L.R., Schulz, R.W., 2013. Thyroid hormone stimulates the proliferation of Sertoli cells and single type A spermatogonia in adult zebrafish (Danio rerio) testis. Endocrinology. 154, 4365–4376. https://doi.org/10.1210/en.2013-1308.

Moussavi, M., Wlasichuk, M., Chang, J.P., Habibi, H.R., 2012. Seasonal effect of GnIH on gonadotrope functions in the pituitary of goldfish. Mol. Cell. Endocrinol. 350, 53–60. https://doi.org/10.1016/j.mce.2011.11.020.

Moussavi, M., Wlasichuk, M., Chang, J.P., Habibi, H.R., 2013. Seasonal effect of gonadotrophin-inhibitory hormone on gonadotrophin-releasing hormone-induced gonadotroph functions in the goldfish pituitary. J. Neuroendocrinol. 25, 506–516. https://doi.org/10.1111/jne.12024.

Moussavi, M., Wlasichuk, M., Chang, J.P., Habibi, H.R., 2014. Seasonal effects of GnIH on basal and GnRH-induced goldfish somatotrope functions. J. Endocrinol. 223, 191–202. https://doi.org/10.1530/joe-14-0441.

Muñoz-Cueto, J.A., Paullada-Salmerón, J.A., Aliaga-Guerrero, M., Cowan, M.E., Parhar, I.S., Ubuka, T., 2017. A journey through the gonadotropin-inhibitory hormone system of fish. Front. Endocrinol. 8, 285. https://doi.org/10.3389/fendo.2017.00285.

Nóbrega, R.H., Greebe, C.D., Van De Kant, H., Bogerd, J., França, L.R., Schulz, R.W., 2010. Spermatogonial stem cell niche and spermatogonial stem cell transplantation in zebrafish. PLoS ONE. 5,12808. https://doi.org/10.1371/journal.pone.0012808.

Paullada-Salmerón, J.A, Cowan, M., Aliaga-Guerrero, M., López-Olmeda, J.F., Mañanós, E.L., Zanuy, S., Muñoz-Cueto, J.A., 2016. Testicular steroidogenesis and locomotor activity are regulated by gonadotropina inhibitory hormone in male European sea bass. PLoS One. 11, 0165494. https://doi.org/10.1371/journal.pone.0165494.

Paullada-Salmerón, J.A., Cowan, M.E., Loentgen, G.H., Aliaga-Guerrero, M., Zanuy, S., Mañanós, E.L., Muñoz-Cueto, J.A., 2019. The gonadotropin-inhibitory hormone system of fish: The case of sea bass (Dicentrarchus labrax). General and Comparative Endocrinology. https://doi.org/10.1016/j.ygcen.2019.03.015.

Schulz, R.W., de França, L.R., Lareyre, J.J., LeGac, F., Chiarini-Garcia, H., Nóbrega, R.H., Miura, T., 2010. Spermatogenesis in fish. Gen. Comp. Endocrinol. 165, 390–411. https://doi.org/10.1016/j.ygcen.2009.02.013.

Sharma, P., Patiño, R., 2013. Regulation of gonadal sex ratios and pubertal development by the thyroid endocrine system in zebrafish. Gen. Comp. Endocrinol. 184, 111–119. https://doi.org/10.1016/j.ygcen.2012.12.018.

Sharma, P., Tang, S., Mayer, G.D., Patiño, R., 2016. Effects of thyroid endocrine manipulation on sex-related gene expression and population sex ratios in zebrafish. Gen. Comp. Endocrinol. 235, 38–47. https://doi.org/10.1016/j.ygcen.2016.05.028.

Swapna, I., Rajasekhar, M., Supriya, A., Raghuveer, K., Sreenivasulu, G., Rasheeda, M.K., Majumdar, K.C., Kagawa, H., Tanaka, H., Dutta-Gupta, A., Senthilkumaran, B., 2006. Thiourea-induced thyroid hormone depletion impairs testicular recrudescence in the air-breathing catfish Clarias gariepinus. Comp. Biochem. Physiol. 144, 1–10. https://doi.org/10.1016/j.cbpa.2006.01.017.

Tovo-Neto, A., da Silva Rodrigues, M., Habibi, H.R., Nóbrega, R.H., 2018. Thyroid hormone actions on male reproductive system of teleost fish. Gen. Comp. Endocrinol. 1, 230–236. https://doi.org/10.1016/j.ygcen.2018.04.023.

Tovo-Neto, A., Martinez, E.R.M., Melo, A.G., Doretto, L.B., Butzge, A.J., Rodrigues, M.S., Nakajima, R.T., Habibi, H.R., Nóbrega, R.H., 2020. Cortisol Directly Stimulates Spermatogonial Differentiation, Meiosis, And Spermiogenesis In Zebrafish (Danio Rerio) Testicular Explants. Biomolecules. 10, 429. https://doi.org/10.3390/biom10030429.

Tsutsui, K., Saigoh, E., Ukena, K., Teranishi, H., Fujisawa, Y., Kikuchi, M., Ishii, S., Sharp, P. J., 2000. A Novel Avian Hypothalamic Peptide Inhibiting Gonadotropin Release. 275, 0–667. https://doi.org/10.1006/bbrc.2000.3350.

Tsutsui, K., Bentley, G.E., Bedecarrats, G., Osugi, T., Ubuka, T., Kriegsfeld, L.J., 2010. Gonadotropin-inhibitory hormone (GnIH) and its control of central and peripheral reproductive function. Front. Neuroendocrinol. 31, 284–295. https://doi.org/10.1016/j.yfrne.2010.03.001.

Tsutsui, K., Ubuka, T., 2013. “Gonadotropin-inhibitory hormone”, in Handbook of Biologically Active Peptides. Section on Brain Peptides, eds A. J. Kastin and H. Vaudry (London: Academic Press). 802–811.

Tsutsui, K., Son, Y. L., Kiyohara, M., Miyata, I., 2017. Discovery of gnih and its role in hypothyroidism-induced delayed puberty. Endocrinol. 159, 62–68. https://doi.org/10.1210/en.2017-00300.

Ubuka, T., Inoue, K., Fukuda, Y., Mizuno, T., Ukena, K., Kriegsfeld, L.J., Tsutsui, K., 2012. Identification, expression, and physiological functions of Siberian hamster gonadotropin-inhibitory hormone. Endocrinology. 153, 373–385. https://doi.org/10.1210/en.2011-1110.

Valle, L.B.S., Oliveira-Filho, R.M., Romaldini, J.H., Lara, P.F., 1985. Pituitary–testicular axis abnormalities in immature male hypothyroid rats. J. Steroid. Biochem. 23, 253–257. https://doi.org/10.1016/0022-4731(85)90402-9.

Zohar, Y., Muñoz-Cueto, J.A., Elizur, A., Kah, O., 2010. Neuroendocrinology of reproduction in teleost fish. Gen. Comp. Endocrinol. 165, 438–455. https://doi.org/10.1016/j.ygcen.2009.04.017.

